# Multiple kinesin-14 family members drive microtubule minus-end-directed transport in plant cells

**DOI:** 10.1101/104471

**Authors:** Moé Yamada, Yohko Tanaka-Takiguchi, Masahito Hayashi, Momoko Nishina, Gohta Goshima

## Abstract

Minus-end-directed cargo transport along microtubules (MTs) is exclusively driven by the molecular motor dynein in a wide variety of cell types. Interestingly, plants have lost the genes encoding dynein during evolution; the MT motors that compensate for dynein function are unknown. Here, we show that two members of the kinesin-14 family drive minus-end-directed transport in plants. Gene knockout analyses of the moss *Physcomitrella patens* revealed that the plant-specific class-VI kinesin-14, KCBP, is required for minus-end-directed transport of the nucleus and chloroplasts. Purified KCBP directly bound to acidic phospholipids (PLs) and unidirectionally transported PL liposomes along MTs *in vitro*. Thus, minus-end-directed transport of membranous cargoes might be driven by their direct interaction with this motor protein. Newly nucleated cytoplasmic MTs represent another known cargo exhibiting minus-end-directed motility, and we identified the conserved class-I kinesin-14 (ATK) as the motor involved. These results suggest that kinesin-14 motors were duplicated and developed as alternative MT-based minus-end-directed transporters in land plants.

## Introduction

Intracellular transport along microtubule (MT) filaments plays a pivotal role in cell organisation and function. Cytoplasmic dynein is a major MT-based motor in animal and fungal cells, and is composed of a motor subunit (dynein heavy chain) and several associated proteins (Cianfrocco et al., 2015; Hancock, 2014; Kardon and Vale, 2009). Upon MT binding, dynein moves processively towards the MT minus end, allowing this motor protein to transport a variety of cargo, from RNA to giant organelles such as the nucleus, along MTs. Interestingly, land plants have lost most of the genes encoding dynein components, including the gene encoding the cytoplasmic dynein heavy chain (Lawrence et al., 2001). How then might plants execute minus-end-directed transport?

It is generally assumed that the actomyosin system is predominantly utilised for cargo transport in plants (Shimmen and Yokota, 2004). However, MT-dependent transport also exists in plants and is shown to be critical in plant physiology in some instances (Kong et al., 2015; Miki et al., 2015; Nakaoka et al., 2015; Zhu et al., 2015). However, little is known about which motors are responsible for transporting which cargoes. In particular, no information is available for minus-end-directed transport.

Two lines of evidence are needed to demonstrate that a specific motor transports a certain intracellular cargo. Firstly, the motor should show processive motility *in vitro*, wherein a single or a cluster of motors take multiple steps unidirectionally along an MT. Dynein, associated with an activator protein, and many, but not all, kinesin family members fulfil this criterion (Cianfrocco et al., 2015; Miki et al., 2005). Secondly, cargo motility and/or distribution within the cell should be perturbed when the motor protein is depleted from the cell. For instance, abnormal mitochondrial distribution in cells depleted of a processive KIF5B (kinesin-1) led to the conclusion that mitochondria are transported by this motor (Tanaka et al., 1998). Similarly, dynein-dependent transport of the nucleus was shown in a filamentous fungus by identifying a dynein mutant with altered nuclear positioning (Xiang et al., 1994). In addition to these two critical data, co-localisation of the motor and cargo constitutes supportive evidence for motor-cargo interaction (Schuster et al., 2011). Another evidence for cargo-motor interaction is *in vitro* reconstitution of the transport process using pure components. For instance, kinesin-3-dependent transport of synaptic vesicles was first shown by observation of kinesin-3 mutant phenotype, followed by *in vitro* reconstitution of liposome transport by purified kinesin-3 (Hall and Hedgecock, 1991; Klopfenstein et al., 2002).

When searching for the minus-end-directed transporter that replaces dynein function in plants, we considered members of the kinesin-14 (hereafter called kin14) family to be candidate motors, because they are known to have minus-end-directed motility along MTs, unlike kinesins from other classes (Endow, 1999). However, the best-studied kin14 in animals, *Drosophila* Ncd, is not processive; Ncd detaches from MTs after only one step along the MT (Case et al., 1997). Land plants have six kin14 subfamilies, five of which are unique to plants (Shen et al., 2012). However, all six kin14 subfamily members are shown to be either non-motile or non-processive in standard single-molecule motility assay (Jonsson et al., 2015; Walter et al., 2015). Interestingly, however, when KCBP-b (class-VI kinesin-14) motors from *P. patens* were artificially tetramerised or tethered on the liposome surface, they moved processively *in vitro* (Jonsson et al., 2015). Furthermore, such processive minus-end-directed motility was observed when Citrine (a GFP variant) was attached to the endogenous KCBP-b motor; punctate Citrine signals representing ≥ 4 Citrine molecules moved 1.0 µm on average before dissociating from MTs (Jonsson et al., 2015). Another study also reported a ~0.18 µm run for KCBP in pavement cells and hypocotyl cells of *Arabidopsis* (Tian et al., 2015). These results suggest that multiple copies of KCBP on cargo surfaces are able to conduct long-distance minus-end-directed cargo transport. Thus, these previous studies provided a candidate minus-end-directed cargo transporter in plants. However, it remained unclear whether KCBP is the motor actually required for minus-end-directed cargo transport in cells.

In the present study, we utilised *in vivo* and reconstitution approaches to test the hypothesis that KCBP directs cargo transport in plants. We focused our *in vivo* observation on the nucleus and newly nucleated MTs in the cytoplasm, which are, to the best of our knowledge, the only two minus-end-directed transport events reported in the plant cytoplasm so far (Miki et al., 2015; Nakaoka et al., 2015). Our data indicated that *P. patens* KCBP is required for minus-end-directed transport of the nucleus immediately after cell division, but not of the newborn MT. We also identified the chloroplast as another KCBP cargo. Furthermore, we reconstituted the direct binding of KCBP to acidic phospholipids and vesicle transport *in vitro*. In addition, we identified class-I kinesin-14 (ATK) as a motor that drives minus-end-directed transport of newborn MTs. Thus, this study identified two kin14s as minus-end-directed transporters in plants for the first time.

## Results

### KCBP is required for minus-end-directed nuclear transport after cell division

To address the role of KCBP in intracellular transport, we used homologous recombination to delete KCBP genes one by one in a line expressing GFP-tubulin and histoneH2B-mRFP. The KCBP subfamily consists of four highly homologous genes in moss (Miki et al., 2014; Shen et al., 2012) and we obtained two independent quadruple knockout (KO) lines (Fig. S1A). The KO line showed cell growth retardation in protonemal tissue, indicating that vigorous moss growth requires KCBP function (Fig. S1B). We assessed the behaviour of the nucleus upon KCBP KO using long-term time-lapse microscopy. Interestingly, we observed a nuclear migration defect upon KCBP KO immediately following cell division in caulonemal apical cells, wherein daughter nuclei did not move towards the cell centre, but stayed near the cell plate (Fig. 1A [0–18 min], 1B, Movie 1). This phenotype was caused by the deletion of KCBP genes, since ectopic expression of KCBP-b rescued the phenotype (Fig. 1B).

**Figure 1.**
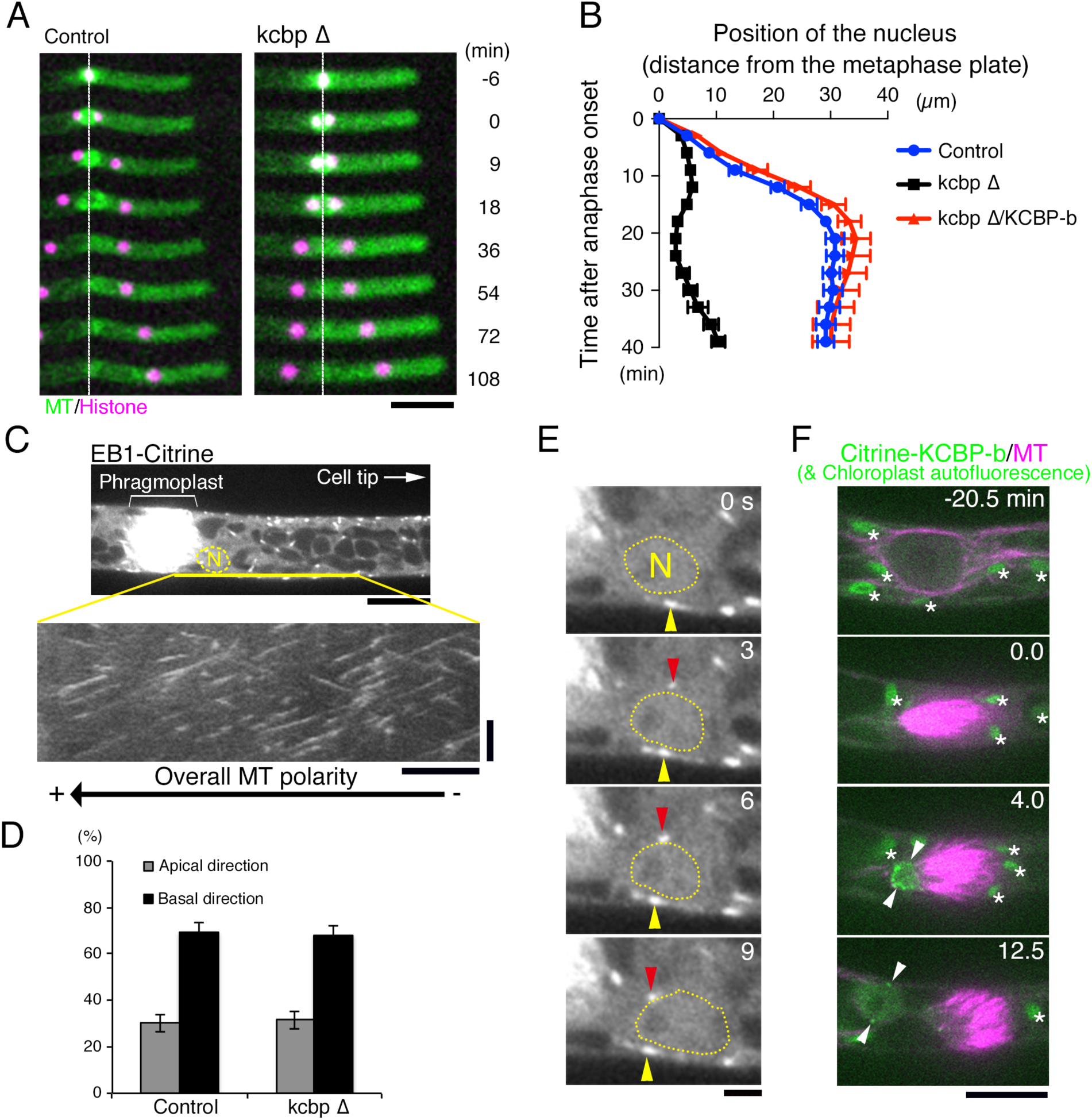
Defective nuclear transport in the absence of KCBP. **(A)** Nuclear migration defect observed immediately following cell division (0–18 min) of a KCBP KO cell. Time 0 corresponds to anaphase onset. Bar, 50 μm. White dot lines indicate the position of the metaphase plate. **(B)** The position of the nucleus after anaphase onset in apical cells. The distance from the metaphase plate is plotted as the mean ± SEM. Control and kcbpΔ, n = 5; kcbpΔ with ectopic KCBP-b expression, n = 6. **(C–E)** MT polarity determination by EB1 imaging. **(C)** Snapshot (top) and kymograph (bottom) of EB1-Citrine, whose moving direction indicates MT polarity. ‘N’ stands for the nucleus. Bar, 10 μm (top). Horizontal bar, 5 μm; Vertical bar, 2 min (kymograph). **(D)** Directionality of EB1 motility was determined using a kymograph that spanned 30 μm from the phragmoplast equator (354 (control) or 248 (KO line) signals in five cells, mean ± SEM). **(E)** EB1-Citrine movement near the nucleus (arrowheads). The tip of this cell is on the right. Bar, 2 μm. **(F)** Localisation of Citrine-tagged KCBP-b proteins. Arrowheads indicate nuclear signals (the other daughter nucleus is out of focus in this movie). Note that autofluorescent chloroplasts are also visible as smaller ellipsoidal structures in this image sequence. Time 0 corresponds to anaphase onset. Bar, 10 μm.

Since nuclear positioning is an MT-dependent process in caulonemal apical cells (Miki et al., 2015), this phenotype in the KO line was attributed to either (1) lack of MT tracks around the nucleus, (2) skewed MT polarity, or (3) defects in minus-end-directed transport. To distinguish these possibilities, we imaged EB1, an MT plus-end-tracking protein, in control and KO lines after cell division (Fig. 1C–E, Movie 2). In both lines, we observed EB1 comets around the nucleus, and the majority of them moved towards the cell plate, not to the cell tip, at this stage (Fig. 1D, E). The data indicated that, regardless of the presence or absence of KCBP, MTs are present around the nucleus and are predominantly oriented with the plus-ends pointed towards the cell plate. Thus, the results suggest that KCBP is required for nuclear transport towards the MT minus-end immediately following cell division.

We analysed KCBP localisation using spinning-disc confocal microscopy in living caulonemal apical cells, in which Citrine was tagged to the amino terminus of the endogenous KCBP-b gene (Fig. 1F, Movie 3). Citrine KCBP-b proteins did not show any particular localisation during metaphase. However, immediately following mitotic chromosome segregation, Citrine signals were transiently enriched near the nuclear region, most likely at the surface of the reforming nucleus (Fig. 1F, arrowheads). The signals appeared 1–2 min after anaphase onset, and were observed for the next 24 ± 5 min (n = 9 ± SD). This duration matched the duration of the KCBP-dependent nuclear movement (Fig. 1B). This finding supports the idea that KCBP is responsible for nuclear transport at this particular stage of the cell cycle.

### KCBP is required for the minus-end-directed chloroplast motility

During cellular observations, we also noticed that chloroplasts were more apically localised in many apical cells upon KCBP KO (Fig. 2A; chloroplasts were visible because of their strong autofluorescence). This notion was confirmed by the kymograph generated over the entire cell cycle (Fig. 2B), as well as by the signal intensity quantification at a fixed time point (Fig. 2C; 150 min after anaphase onset). Chloroplast positioning depends on both actin and MTs (Sato et al., 2001). To test which cytoskeletal filament is required for chloroplast positioning in our conditions, we depolymerised F-actin or MTs by treating caulonemal cells with specific inhibitors (Fig. S2). We observed a profound impact on chloroplast positioning after MT destabilisation by oryzalin, whereas no clear positioning defect was observed after actin destabilisation by latrunculin A. We concluded that MTs are critical for uniform distribution of the chloroplasts.

**Figure 2.**
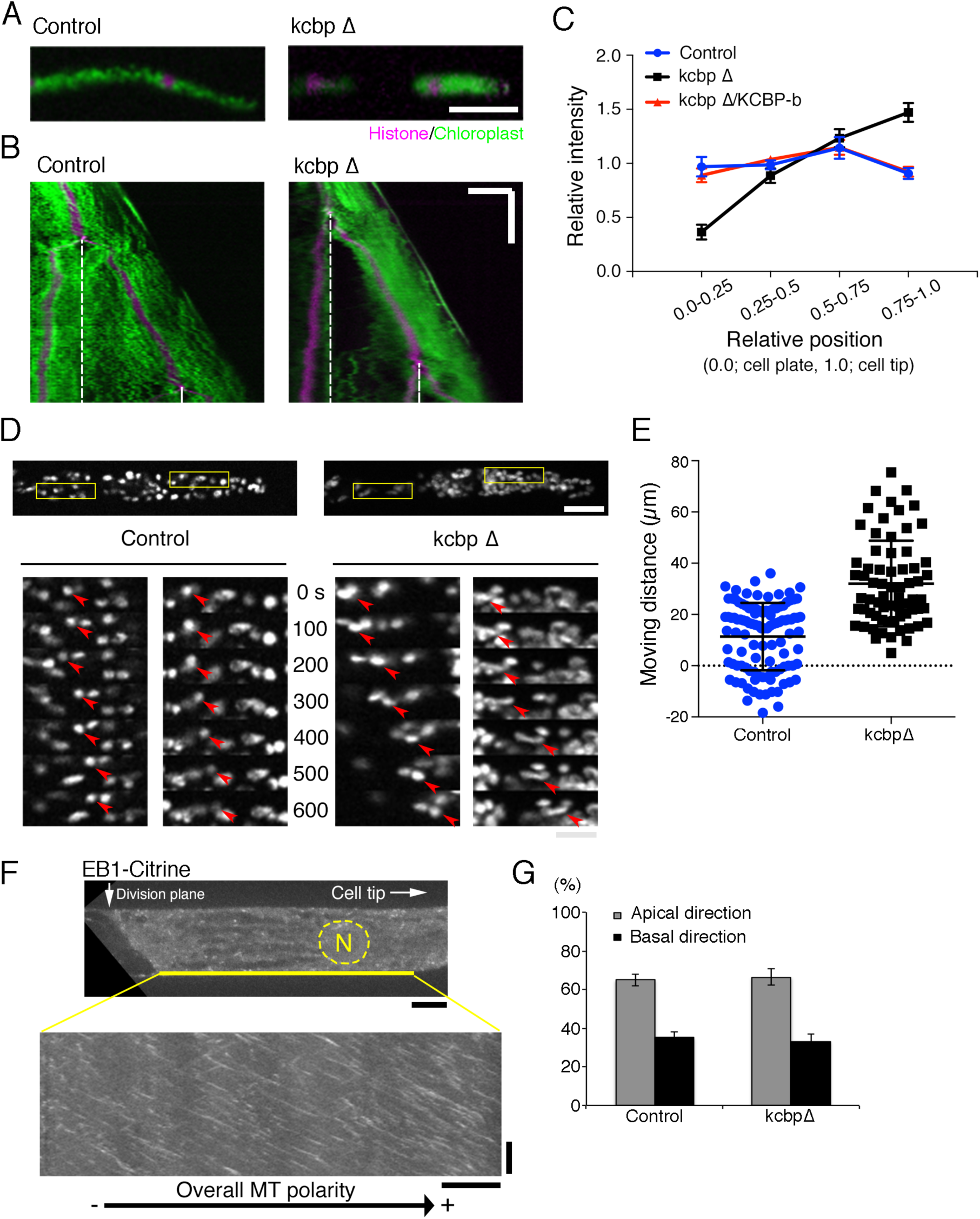
Abnormal chloroplast ditribution in KCBP KO cells. **(A)** Snapshots of KCBP KO cells at 150 min after anaphase onset. Green; chloroplast, Magenta; histone-RFP. Bar, 50 μm. **(B)** Kymograph showing chloroplast distribution along the apical-basal axis throughout the cell cycle. White dotted lines indicate the position of the metaphase plate. **(C)** Chloroplast signal intensity was measured along cells’ long axes. Each cell was divided into four regions, and the mean signal intensity within each region was divided by the average intensity of the whole cell. The mean relative values and SEMs are shown in this graph. Control, n = 8; knockout line, n = 14. **(D)** Apical translocation of chloroplasts in KCBP KO. Arrowheads indicate tracking of two chloroplasts. Bars, 20 (top) and 10 (bottom) μm. **(E)** Net displacement of individual chloroplasts along cell’s long axis in 30 min. Negative values were given when the chloroplast moved towards cell plate (basal direction). Mean and SD are marked (p < 0.0001; Mann-Whitney U-test). **(F–G)** Directionality of EB1 signal motility during 60–90 min after anaphase onset (407 (control) or 235 (KO line) signals in five cells, mean ± SEM). A snapshot and kymograph are displayed. Bar, 10 μm (top). Horizontal bar, 10 μm; Vertical bar, 2 min (kymograph).

To confirm whether the abnormal distribution of chloroplasts in the KCBP KO line is caused by defects in their minus-end-directed transport, we aimed to observe chloroplast dynamics at shorter time intervals (10 s). However, chloroplasts were apically enriched in the KO line in majority of the cell cycle, which hindered individual chloroplast tracking. Therefore, we focused on post-anaphase apical cells (~60 min after anaphase onset), in which new cell plate deposition near the nucleus reset chloroplast distribution within the cell; several individual chloroplasts were traceable for > 30 min, regardless of the presence or absence of KCBP (Fig. 2D, 2E, Movie 4). Our manual tracking of 108 chloroplasts in the control line indicated that chloroplasts moved back and forth, and net displacement was modest after 30 min; on average, they moved apically at a velocity similar to the tip growth speed (Fig. 2E, S1B). In sharp contrast, in the KO line, chloroplasts tended to translocate more apically, with suppression of basal motility (n = 68). This observation explained the apical accumulation of chloroplasts in the KO line in majority of the duration of cell cycle. At this stage, more than 60% of the EB1 signals moved towards the apical cell tip, suggesting that MTs are overall oriented in such a manner that plus-ends are facing the cell tip (Fig. 2F–G; (Hiwatashi et al., 2014)). These results are consistent with the idea that, in the absence of KCBP-dependent minus-end-directed motility, one or more counteracting plus-end-directed motors move chloroplasts to the tip. Thus, the minus-end-directed transport defect in the absence of KCBP was not limited to the nucleus.

### *In vitro* reconstitution of KCBP-dependent vesicle transport

KCBP has the FERM (4.1/ezrin/radixin/moesin) domain in its tail region, which is known to bind lipids (Fig. 3A) (Chishti et al., 1998). Given the identification of two membranous cargoes in cells, an interesting possibility was that KCBP might bind to membranes directly. However, in our previous study, we failed to observe direct binding of purified KCBP to a liposome made with phospholipids (PLs) even after several attempts; therefore, we artificially tethered purified histidine-tagged KCBP-b to the liposome surface by introducing a Ni-NTA-conjugated lipid, and observed liposome motility *in vitro* (Jonsson et al., 2015). In the present study, motivated by the defects in organelle distribution in the KCBP KO line, we revisited the liposome-binding assay for this motor.

**Figure 3.**
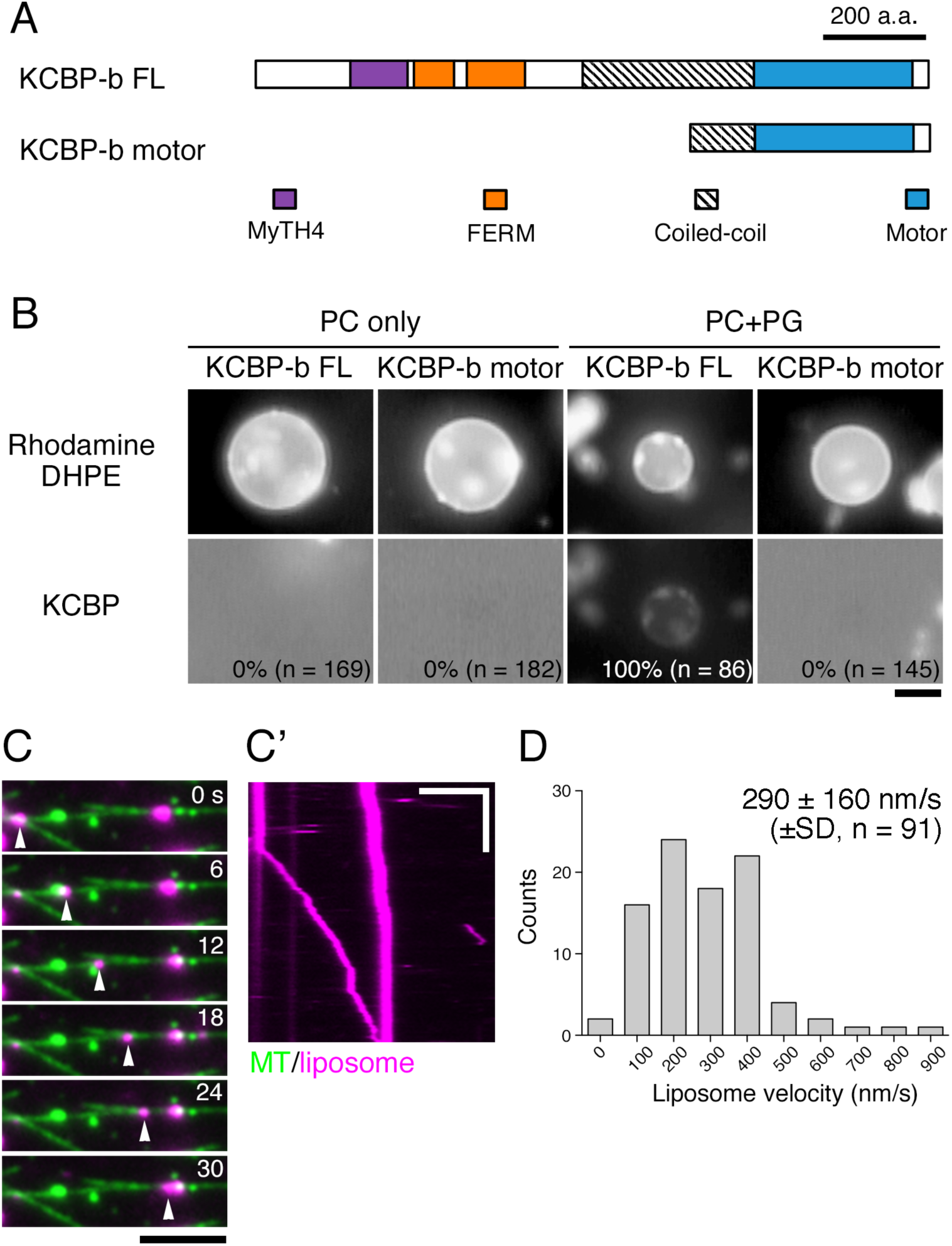
Reconstitution of liposome transport by KCBP. **(A)** Domain organisation of full-length (FL) and truncated KCBP-b proteins (851–1322 aa) used in the in vitro assay. **(B)** Reconstitution of KCBP-b liposome binding. Full-length GFP-KCBP-b bound to the liposome composed of PC and PG, but not the liposome made solely with PC. Full-length GFP-KCBP-b, but not truncated protein, bound to liposomes. Frequency of GFP-positive liposomes is indicated in each panel. Background GFP signals are high in three panels because of diffused GFP-kinesins. Bar, 5 μm. **(C, D)** Reconstitution of liposome transport by KCBP-b. Image sequences, kymograph, and velocity distribution are shown. Arrowheads indicate a fast-moving smaller liposome, whereas a larger liposome is moving slowly. Horizontal bars, 2 μm; vertical bar, 10 s.

In the previous study, we used liposomes made dominantly (90%) of phosphatidylcholine (PC), a neutral phospholipid, although the PC content of cellular membranes is typically < 50% (Dewey and Barr, 1971; Novitskaya et al., 2000; Schwertner and Biale, 1973; Whitman and Travis, 1985). To better mimic cellular membranes in terms of composition of neutral and acidic PLs, we prepared giant hybrid liposomes composed of PC (50%) and an acidic phospholipid (50%; phosphatidylglycerol (PG), phosphatidylamine (PA), or phosphatidylserine (PS)). Interestingly, we observed that purified GFP-KCBP-b decorated all three hybrid liposomes, but not those assembled solely with PC (Fig. 3B; PC only and PC-PG liposome are displayed). This result explained why we failed to observe direct binding in the previous study. Liposomal binding of KCBP was tail-dependent, since we did not observe GFP decoration around the PC-PG liposomes when a construct lacking a tail was used (Fig. 3B). These results indicated that KCBP-b is capable of directly binding to acidic phospholipids. Furthermore, it suggests that KCBP has an ability to bind to many, if not all, membranes in the cell.

To address if KCBP can transport the PL liposomes along MTs, we prepared full-length GFP-KCBP-b and smaller PC-PG liposomes that were fluorescently labelled. When they were supplied to flow chambers with MTs attached to a coverslip, unidirectional transport of the liposomes along MTs was observed (Fig. 3C, C′, D, Movie 5). These reconstitution results demonstrated the ability of the KCBP-b motor to transport membrane vesicles via direct binding.

### ATK is required for minus-end-directed MT transport

Another cargo that has been demonstrated to be transported in the minus-end direction in plant cells is the MT that is nucleated in a branching fashion in the cytoplasm (Nakaoka et al., 2015). Since MT association with the tail domain of *Arabidopsis* KCBP was previously shown (Tian et al., 2015), we investigated if moss KCBP is required for branching migration of MTs. However, we observed branching MT migration in the absence of all four KCBPs (Fig. 4A). The result indicates that yet another minus-end-directed motor is involved in MT transport.

**Figure 4.**
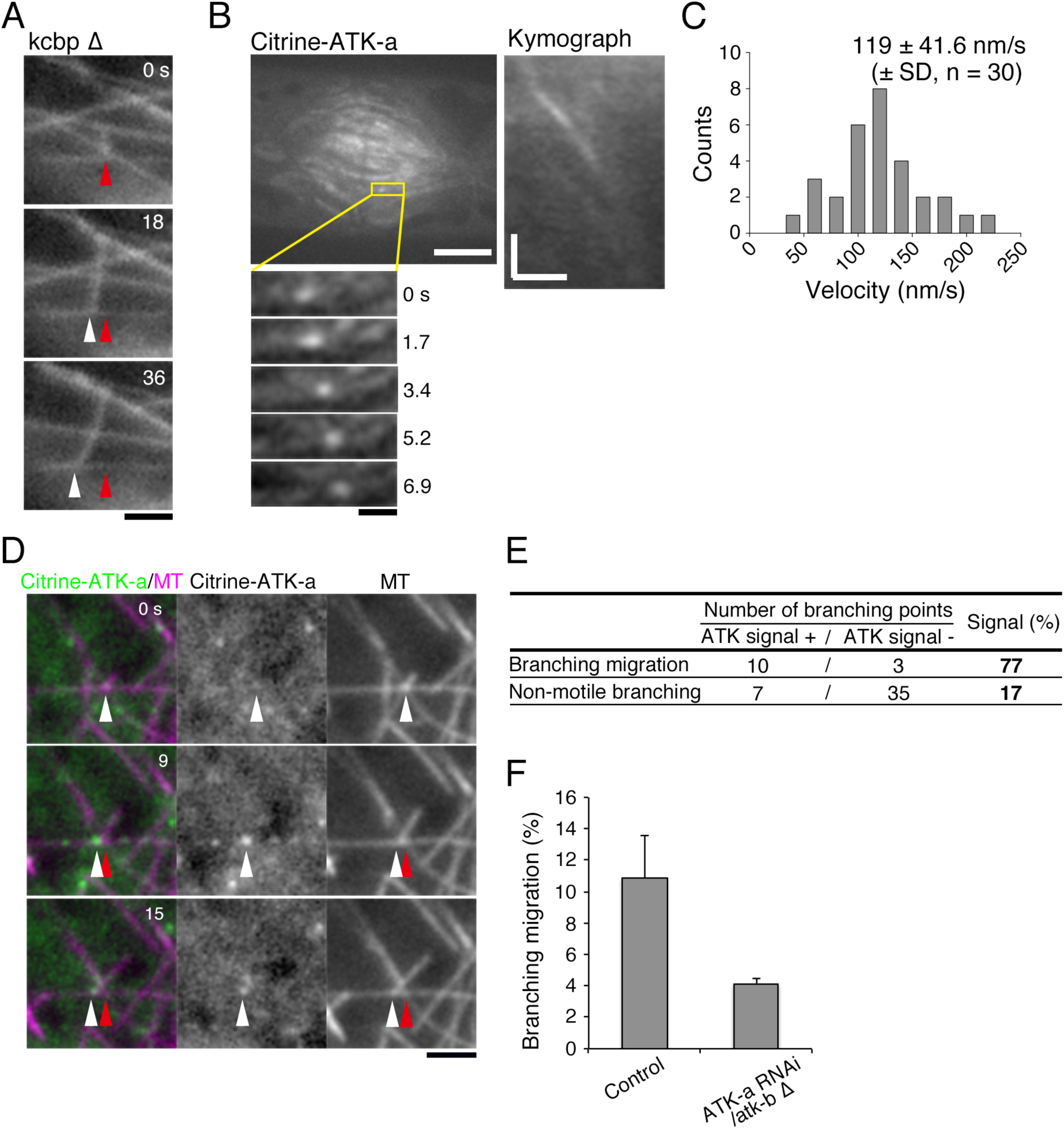
ATK depletion reduces branching MT migration. **(A)** Branching MT migration observed in the quadruple KCBP KO line. Bar, 2 μm. **(B)** Poleward movement of Citrine-ATK-a signals in the phragmoplast. Image sequences and kymographs are shown. Bars, 5 μm (top), 1 μm (bottom left), and 10 μm/2 sec (kymograph). **(C)** The velocity of Citrine-ATK-a motility in the phragmoplast. **(D)** Citrine-ATK-a signal detected at the migrating branch points (arrowheads). Bar, 2 μm. **(E)** Frequency of Citrine-ATK-a localisation at the migrating or non-migrating branching point. **(F)** Frequency of branching MT migration (per branching nucleation, ± SE) (n = 2 [control ATKbΔ] or 3 experiments [ATK-a/-IbΔ RNAi]). 21–37 branching MTs were identified in 3 cells for each experiment.

We reasoned that another kin14 family member plays such a role. The ATK/kin14-I subfamily is the only subfamily of kin14 for which orthologues are found in animals (*P. patens* ATK-a/ATK-b, *Arabidopsis* ATK1/ATK5, *Drosophila* Ncd, human HSET). Although the dimeric Ncd motor is non-processive, a previous study showed that artificial clustering of the Ncd motor enabled processive minus-end-directed motility *in vitro* (Furuta et al., 2013). Similar to results observed for animal orthologues, Citrine-ATK-a was predominantly localised in the nucleus during interphase and decorated MTs during mitosis (Fig. S3A). Interestingly, spinning-disc confocal microscopy occasionally revealed punctate Citrine-ATK-a signals that moved polewards at a velocity of ~120 nm/s (n = 30) in the phragmoplast (late mitotic apparatus) (Fig. 4B, C, Movie 6 [right]). Furthermore, surprisingly, when oblique illumination fluorescence microscopy that excludes dominant nuclear signals from the imaging field was used, we observed Citrine-ATK-a spots transiently occurring on cytoplasmic MTs, and some of those spots coincided with MT branching points, where a ‘daughter’ MT was migrating along a ‘mother’ MT (Citrine signals were detected in 10 of 13 migrating MTs; Fig. 4D, E, Movie 6 [left]). MT migration correlated closely with ATK localisation, as the signals were detected at a much lower frequency (17% [n = 42]) on non-migrating MTs that were nucleated in a branching fashion (Fig. 4E). Thus, ATK has become a candidate minus-end-directed transporter for branching MTs.

We next aimed to test if ATK depletion affects the frequency of branching MT migration in the cytoplasm. However, we could not obtain a double ATK-a and ATK-b KO line, even after multiple attempts, although ATK-a or ATK-b single KO lines grew normally. We reasoned that these proteins redundantly execute an essential function in moss. Therefore, to assess the function of ATK, we selected conditional RNAi lines (Nakaoka et al., 2012), in which an inducible RNAi construct targeting the ATK-a gene was integrated into the genome of the ATK-b KO line. Upon RNAi induction in this line, we frequently observed frayed mitotic spindles and phragmoplasts (Fig. S3B, C). The phenotype was consistent with the ATK family in other cell types that undergo acentrosomal cell division (Ambrose and Cyr, 2007; Chen et al., 2002; Hatsumi and Endow, 1992; Ito and Goshima, 2015). The phenotype was rescued by the expression of RNAi-insensitive ATK-a gene (Fig. S3C).

Using the established RNAi line, we observed the effects of ATK depletion on branching MT migration in the cytoplasm using oblique illumination fluorescence microscopy. Consistent with our previous report (Nakaoka et al., 2015), we occasionally observed MT nucleation from the lattice of an existing MT, among which ~10% of the nucleated daughter MTs moved along mother MTs. However, branching migration frequency was reduced following ATK-a RNAi (Fig. 4F). These results indicate that ATK is required for minus-end-directed transport of branching MTs in the cytoplasm.

## Discussion

In this study, we identified two kin14 motors as MT-based minus-end-directed transporters in moss. To the best of our knowledge, these are the first cytoplasmic minus-end-directed transporters identified in plants that have lost the cytoplasmic dynein complex during evolution. Among three cargoes identified, MTs and the nucleus are well known cargoes of dynein in animals (Rusan et al., 2002; Tanenbaum et al., 2010). Minus-end-directed transport has been underappreciated in plants because of the lack of dynein and the predominance of the actomyosin system for intracellular dynamics in traditional model cell types (Shimmen and Yokota, 2004; Vale, 2003). In fact, the loss of dynein in plants appeared to be partially compensated for by the development of the actomyosin system; for instance, the actin-based motor myosin XI-i is critical for nuclear motility in a few cell types in *Arabidopsis*, and is essential for moss viability (Tamura et al., 2013; Vidali et al., 2010). In addition, we propose that plants developed an alternative motor-based system, wherein kin14 motors were duplicated and utilised as MT-based minus-end-directed transporters (Fig. 5). Consistent with this notion, a previous study identified MTs, not actin, as the critical cytoskeletal filament for nuclear migration in tobacco microspores (Zonia et al., 1999). It is of interest to investigate how actomyosin- and kinesin/MT-dependent mechanisms are differentially and/or cooperatively utilised in various plant cell types.

**Figure 5.**
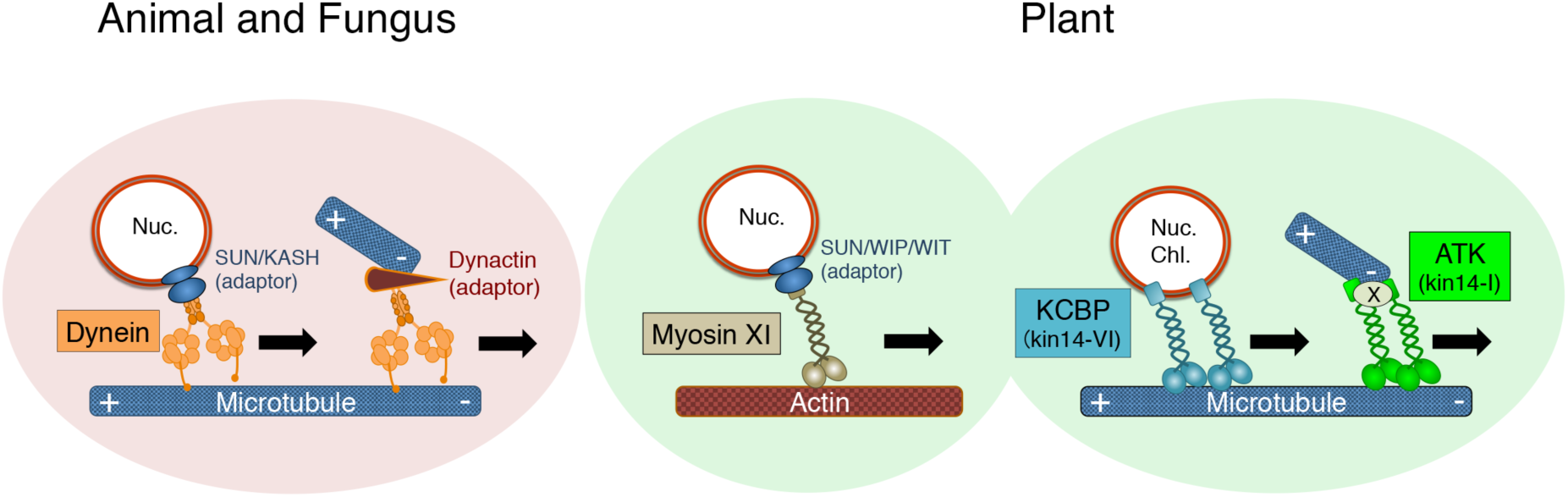
Long-distance transport in plants and animals. In animal cells, dynein is considered to be the sole transporter for long-distance, minus-end-directed transport along MTs. However, land plants have lost the dynein gene. Plants have developed an actin-based transport system that utilises plant-specific myosin-XI (Ueda et al., 2015), as well as the MT- and kinl4-dependent mechanisms (revealed in this study). Unlike dynein, at least two different kinl4s are involved in the minus-end-directed transport mechanism: kinl4-VI/KCBP transports the nucleus, chloroplasts, and possibly other membranous cargoes via direct binding, whereas kinl4-I/ATK drives MT transport along other MTs. ATK clusters at the migrating point, which might require an additional factor (depicted as factor X in this model).

### KCBP is likely a versatile minus-end-directed transporter

Several results of this study indicated that KCBP is a minus-end-directed cargo transporter. First, minus-end-directed motility of the nucleus and chloroplasts are suppressed upon KCBP deletion. Although loss of properly polarised MTs would lead to similar consequences, our EB1 imaging did not support this alternative possibility. Second, KCBP was enriched at the nuclear surface during the portion of the cell cycle where the nucleus exhibits minus-end-directed motility. It is expected that a large cargo like the nucleus would require the action of multiple motors for long-distance delivery (Tanenbaum et al., 2010). Whether KCBP also accumulated at the chloroplast surface remains unclear because of overwhelming autofluorescence of this organelle. Finally, PL liposomes were transported *in vitro* by purified KCBP protein without the aid of artificial crosslinkers or any other proteins. The result suggests that KCBP cargo might not be limited to these two organelles; virtually every membranous material, from small vesicles to giant organelles, is a cargo candidate for KCBP. In addition, the tail domain of KCBP binds to MTs and actins (Tian et al., 2015); it is possible that these cytoskeletal filaments and also other proteins are transported by KCBP through direct binding or through adaptor molecules. Thus, growth retardation in moss (Fig. S1B) or abnormal trichome morphology in *Arabidopsis* (Oppenheimer et al., 1997; Tian et al., 2015) upon KCBP KO might result from defects in transport of various cargoes.

Our data also suggests that KCBP activity is regulated in a cell cycle-dependent manner. We observed transient accumulation of KCBP around the nucleus, and nuclear migration defects were observed only during this period upon KCBP KO. The mechanism of KCBP regulation during the cell cycle is currently unclear. Despite the initial migration defect in the absence of KCBP, the nucleus eventually moved towards the cell centre prior to mitosis, suggesting that redundant mechanisms are also present in moss cells (Fig. 1A, 36–108 min). *In vitro* studies indicated that two other kin14 family members, kin14-II and kin14-IV, have minus-end-directed motility, albeit non-processive, and these motors are also candidate minus-end-directed transporters in cells (Jonsson et al., 2015; Walter et al., 2015). Interestingly, nuclear binding of and transport by kinesin and dynein is a regulated process in animals as well, whose molecular mechanisms are largely unclear (Tanenbaum et al., 2010; Tsai et al., 2010).

### ATK is a minus-end-directed MT transporter

ATK also fulfils the two criteria for a cargo transporter. First, ATK depletion from moss cells reduced the frequency of branching MT migration, namely, minus-end-directed transport of newly nucleated MTs. We speculate that incomplete elimination of this event in the RNAi line is due to the presence of residual ATK protein, although it cannot be ruled out that other motor proteins are also involved in this process. Second, ATK was often concentrated at the branching point during migration in cells. Notably, the observed branching migration was reminiscent of dynein-dependent inward transport of cytoplasmic MTs during the mitotic reorganisation of MT arrays in animal cells (Rusan et al., 2002). Our results suggest that plant ATK has acquired the MT-transporting activity that dynein possesses in animals.

It is unlikely that overall MT organisation during interphase is grossly affected by ATK depletion, since branching MT nucleation accompanying with migration constitutes only 3% of the total MT generation events in this cell type; more dominant modes are cytoplasmic nucleation in which MTs are nucleated spontaneously in the cytoplasm without template mother MTs (56%), branching MT nucleation that does not accompany migration (30%), and MT severing (11%) (Nakaoka et al., 2015). Loss of ATK, however, had a profound effect on spindle/phragmoplast coalescence. This can be partly attributed to the loss of MT bundling and sliding activities of this motor, as has been proposed for its animal counterparts (Fink et al., 2009). However, given that we detected a cluster of GFP-ATK-a moving poleward, it is tempting to speculate that ATK-dependent branching migration takes place in the spindle/phragmoplast and assists in MT coalescence. Since MTs were crowded, particularly in the metaphase spindle, branching MT migration or minus-ends of MTs could not be directly observed. Nevertheless, in support of this idea, a previous study using tobacco BY-2 cells reported poleward movement of the putative minus-end marker γ-tubulin in the phragmoplast (Murata et al., 2013). Furthermore, in ATK-dependent branching migration observed in the interphase cytoplasm, the angle between daughter and mother MTs becomes shallower after migration (Nakaoka et al., 2015), suggesting that this mode of crosslinking is an excellent means to align two MTs in a parallel manner as well in the spindle/phragmoplast. Whether the ATK motor can self-cluster at MT branching points or it requires additional factors, such as those that induce multimerisation, remains to be determined.

## Materials and methods

### Moss culture, plasmid construction, gene disruption, and Citrine tagging

Plasmids and primers used for protein expression, gene disruption, genotyping, and Citrine tagging are listed in Table S1. Moss lines are listed in Table S2. Methodologies of moss culture, transformation, and transgenic line selection (Citrine tagging) were previously described thoroughly (Yamada et al., 2016). Briefly, we used BCDAT medium for routine culture. Transformation was performed by the standard PEG-mediated method. Citrine tags were added to the N-termini of kin14 genes via homologous recombination (drug-resistant genes were not integrated into the genome). Gene knockouts were obtained by replacing an endogenous kin14 gene with a drug-resistant marker flanked by lox-P sequences. When two genes were depleted, two marker cassettes were sequentially integrated into each gene locus. For triple or quadruple knockout selection, the existing two markers were removed by transient expression of Cre recombinase, followed by replacing the other two genes with the same markers (Cre-expressing plasmid pTN75 was a gift from Dr. Mitsuyasu Hasebe [NIBB, Japan]). Gene disruption was confirmed by PCR.

### In vivo microscopy

The methodology for inducible RNAi was previously described thoroughly (Miki et al., 2016). For mitosis imaging after conditional ATK-a RNAi, cells were pre-cultured in BCD medium for 4 days, followed by RNAi induction with 1 µM β-estradiol for 3 days. Branching migration was quantified after 10 days of RNAi induction (RNAi was less penetrant in this condition because cells had to be plated on a cellophane-coated medium for oblique illumination microscopy). Oryzalin (10 µM; AccuStandard), latrunculin A (25 µM; Wako), or control DMSO (1%) was added to protonemal cells plated on culture medium for 5–6 days. Methods for epifluorescence and spinning-disc confocal microscopy were previously described thoroughly (Yamada et al., 2016). Briefly, protonemal cells were plated onto glass-bottom plates coated with BCD agar medium. Long-term imaging by a wide-field microscope (low magnification lens) was performed with a Nikon Ti (10X 0.45 NA lens, EMCCD camera Evolve [Roper]). High-resolution imaging was performed with a spinning-disc confocal microscope (Nikon TE2000 or Ti; 100X 1.40 NA lens, 100X 1.45 NA lens or 20X 0.75 NA lens, CSU-X1 [Yokogawa], EMCCD camera ImagEM [Hamamatsu]). Oblique illumination microscopy was performed as previously described (Jonsson et al., 2015; Nakaoka et al., 2015); cells were cultured in BCDAT medium and a Nikon Ti microscope with TIRF unit, a 100X 1.49 NA lens, GEMINI split view (Hamamatsu), and EMCCD camera Evolve (Roper) was used. Branching migration was also scored using this microscopy setup. Images were acquired every 3 s for 10 min. All imaging was performed at 24–25°C under dark condition except for growth speed imaging, during which white light was illuminated for 100 s between image acquisitions. Microscopes were controlled by micromanager.

### Protein purification

KCBP-b proteins used in this study have identical sequences to those previously reported (Jonsson et al., 2015). His-mGFP-KCBP-b (full-length) proteins were purified with Ni-NTA beads from insect Sf21 cells, where the lysis buffer contained 25 mM MOPS (pH 7.0), 2 mM MgCl_2_, 250 mM NaCl, 5% sucrose, 5 mM β-mercaptoethanol, 1 mM ATP, 30 mM imidazole, 1% Triton X-100 and protease inhibitors (0.5 mM PMSF and peptide cocktails [1 µg/ml leupeptin, pepstatin, chymostatin, and aprotinin]), and the elution buffer contained 25 mM MOPS (pH 7.0), 2 mM MgCl_2_, 250 mM NaCl, 5% sucrose, 5 mM β-mercaptoethanol, 1 mM ATP and 400 mM imidazole. Truncated His-GFP-KCBP-b (851–1322 aa) was expressed in *E. coli* BL21-AI with 0.2 % arabinose and 500 µM IPTG for 18 h at 18°C. Harvested cells were lysed with the Advanced Digital Sonifier D450 (Branson) in the lysis buffer described above, followed by purification using Ni-NTA beads. Imidazole was removed at the final step by dialysis using PD MiniTrap G-25 column (GE Healthcare) with the elution buffer. Proteins were flash frozen and stored at −80°C.

### In vitro liposome binding/motility assay

To prepare and observe giant liposomes, a mixture of phospholipids, egg (Powder) phosphatidylcholine (PC; Avanti), egg (Powder) phosphatidylglycerol (PG; Avanti), and rhodamine DHPE (Molecular Probes) (50:50:0.3, mol/mol), was dissolved in chloroform, dried under a constant stream of N_2_ gas, and desiccated in a vacuum for at least 2 h to produce dried lipid film (Tanaka-Takiguchi et al., 2013). This film was then hydrated with 420 mM sucrose at 50°C for 120 min to obtain a liposome suspension. For the kinesin-liposome binding assay, the liposome suspension was diluted 20-fold with 1X assay buffer (25 mM MOPS (pH 7.0), 2 mM MgCl_2_, 75 mM KCl and 1mM EGTA) (final 40 µM lipids), mixed with each kinesin sample (400 nM), and then observed by fluorescence microscopy (BX60 microscope [Olympus] with 100X 0.75 NA lens and WAT-910HX CCD camera [Watec]). Transport assays were performed by a previously described motility assay protocol (Jonsson et al., 2015) with some modifications. Importantly, KCBP protein was directly bound to liposomes in the current study, whereas in the previous study, histidine-tagged KCBP was bound to liposomes that contained Ni-NTA-conjugated lipids (Jonsson et al., 2015). Anti-biotin (1–5% in 1X MRB80; Invitrogen) was added to a flow chamber made with silanized coverslips and allowed to incubate for 5 min. The flow chamber was washed with 1X assay buffer and supplied with 1% Pluronic F127 (another 2–3 min incubation). The flow chamber was then washed with the assay buffer, followed by 5 min incubation with labelled MTs (10% Cy5-labelled tubulin and 10% biotin-labelled tubulin) and 40 µM taxol. The flow chamber was washed with the assay buffer that contained 40 µM taxol. Finally, kinesin motors in the assay buffer (with 2 mM ATP, 20 µM taxol, 0.1% methylcellulose, 0.5 mg/ml casein and oxygen scavenger system (50 mM glucose, 400 µg/ml glucose-oxidase, 200 µg/ml catalase and 4 mM DTT)) and 1 µM liposomes were added. Liposomes and kinesin solutions were mixed immediately prior to imaging. The liposome solution was extruded through a polycarbonate filter with 200 nm pore size using a mini extruder (Avanti) and the motors were subjected to MT binding-and-release to select active motors.

### Data analysis

To quantify the cell growth rate, we acquired cell images every 3 min for 10 h. Kymographs were then generated for tip-growing caulonemal cells by using ImageJ and the tip growth rate was measured. In some cases, growth velocity decreased from the middle of the image sequences, perhaps due to shortage of light-dependent energy; in such cases, initial growth speed was measured. Quantification of the nuclear movement following cell division was performed for caulonemal apical cells. Images were acquired every 3 min for 10 h under dark condition and kymographs were generated. The distance from the metaphase plate to the nuclear centre was measured after generating kymographs. To analyse MT polarity in post-anaphase cells, EB1-Citrine or EB1-mCherry was imaged every 3 s using the spinning-disc confocal microscope (Kosetsu et al., 2013). Directionality of the EB1 motility was determined after generating kymographs along cell’s long axis, which spanned 30 µm from the cell plate. MT polarity in interphase was determined in a similar way. The velocity of Citrine-ATK-a motility in the phragmoplast was quantified also based on kymographs. Images were acquired at the phragmoplast surface every 0.8 s. Citrine-ATK-a/atk-b ∆ line was used for this assay. ATK localisation during interphase was analysed based on images acquired every 3 s with oblique illumination fluorescence microscopy. In analysing ATK localisation at non-motile branching points, we focused on the branch at which daughter MT length was less than 3 µm, since we observed that 8 of 9 migrating daughter MTs were < 3 µm in length. To quantify chloroplast distribution, the intensitiy of chloroplast autofluorescence was measured (visualized with 640 nm laser). Images were acquired every 3 min for 10 h under dark condition and the chloroplast signal intensity was measured along cells’ long axes by drawing a one-pixel-wide line using ImageJ. Each cell was divided into four regions, and the mean signal intensity within each region was divided by the average intensity of the whole cell. For the liposome-kinesin binding experiment, we randomly selected liposomes with a diameter of more than 2 µm and checked if the liposome had ring-shaped GFP signal, which represented membrane association. When several liposomes formed an aggregate, we counted the aggregate as one liposome.

### Accession numbers

Sequence data used in this article can be found in the Phytozome database by following accession numbers. *KCBP-a*, Pp3c15_3730; *KCBP-b*, Pp3c9_4530; *KCBP-c*, Pp3c2_29920; *KCBP-d*, Pp3c11_5890; *ATK-a*, Pp3c7_11530; *ATK-b*, Pp3c11_17850.

## Acknowledgements

We thank Mitsuyasu Hasebe and Mamoru Sugita for plasmids; Rie Inaba and Yuki Nakaoka for technical assistance; Tomohiro Miki for helpful comments on the manuscript; Yuichiro Maéda for generous support; and Kingo Takiguchi for the helpful discussion. This work was funded by the TORAY Science Foundation and JSPS KAKENHI 15K14540 (to G.G). M.Y. is a recipient of a JSPS pre-doctoral fellowship. The authors declare no competing financial interests.

## Supplemental Movie Legends

**Movie 1. Nuclear migration immediately following cell division is defective in the quadruple KCBP KO line**

Images were acquired every 30 s with spinning-disc confocal microscopy. Green; MT, magenta; histone. Time 0 corresponds to the timing of nuclear envelope breakdown.

**Movie 2. EB1-Citrine imaging during telophase**

Images were acquired every 3 s with spinning-disc confocal microscopy. The nucleus of the tip cell is indicated by arrowhead.

**Movie 3. Localisation of Citrine-KCBP-b**

Citrine-KCBP-b (green) and MT (magenta) were imaged every 30 s with spinning-disc confocal microscopy. Arrowheads indicate Citrine-KCBP-b localisation around the nucleus (the other daughter nucleus is out of focus). Note that autofluorescent chloroplasts are also visible in the green channel. Time 0 corresponds to the timing of anaphase onset.

**Movie 4. Chloroplast movement is defective in KCBP KO**

Chloroplasts of post-anaphase apical cells (~60 min after anaphase onset) were imaged every 10 s with spinning-disc confocal microscopy.

**Movie 5. Liposome transport by KCBP *in vitro***

Unidirectional motility of liposomes (magenta) along MTs (green) in the presence of full-length Citrine-KCBP-b. Liposome images were acquired every 0.5 s with total internal reflection fluorescence microscopy. MTs were imaged at a single time point prior to liposome imaging.

**Movie 6. Motility of Citrine-ATK-a signals during branching MT migration and in the phragmoplast**

Oblique illumination fluorescence microscopy (left) and spinning-disc confocal microscopy (right). Green; Citrine-ATK-a, magenta; MT. Arrowheads indicate moving Citrine-ATK-a signals.

**Figure S1.**
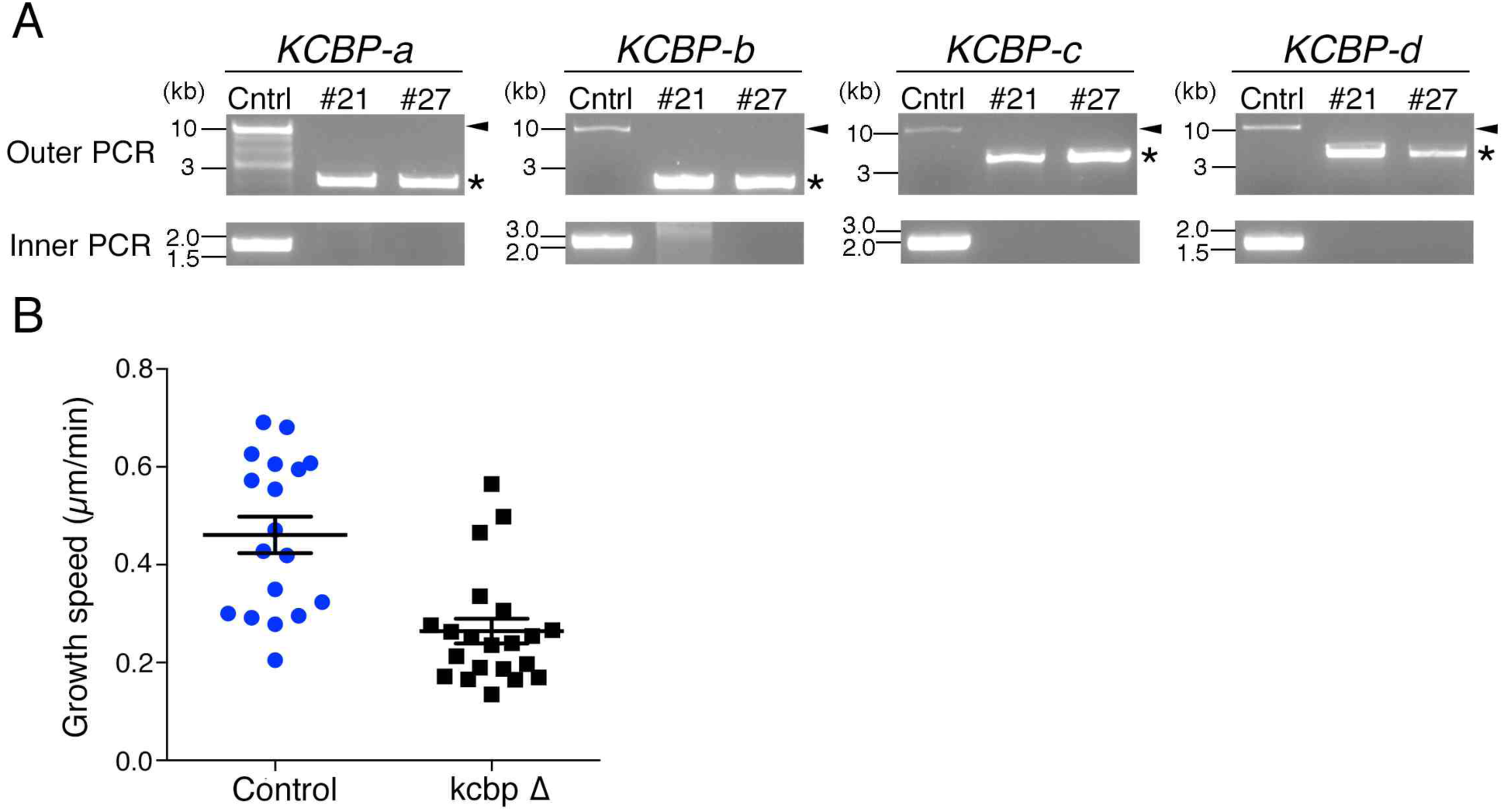
Growth retardation by deleting all four KCBP genes. **(A)** Confirmation of the quadruple KCBP KO line by PCR. Genomic DNAs from a control line and two KCBP KO lines (#21 and #27) were ued as PCR templates. ‘Outer’ PCR primers were designed at 5′ and 3′UTR of each gene, whereas ‘Inner’ pimers targetted the open reading frame. Expected size in control or the KO line was marked by arrowheads or asterisks, respectively. **(B)** Slower tip growth of the caulonemal apical cell in the KCBP KO line. Control, n = 18; KO line, n = 21, ± SE, p < 0.0001 (Mann-Whitney U-test).

**Figure S2.**
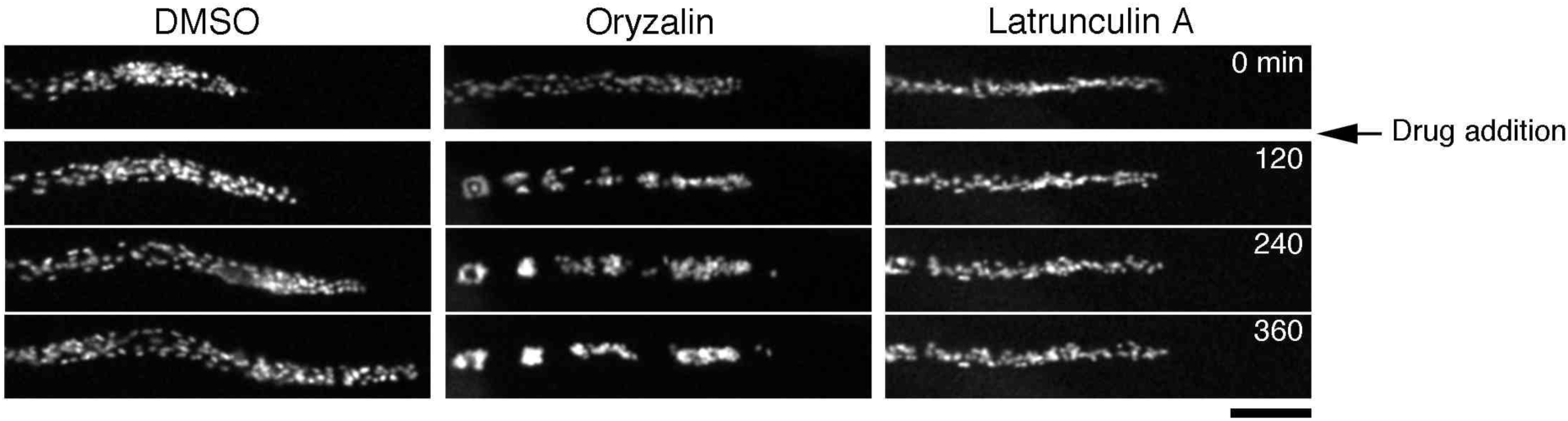
Chloroplast positioning requires MT cytoskeleton. Chloroplast positioning after MT or actin destabilisation. Oryzalin (10 μm), latrunculin A (25 μM), or control DMSO (1%) was added at time 0. Bar, 50 μm.

**Figure S3.**
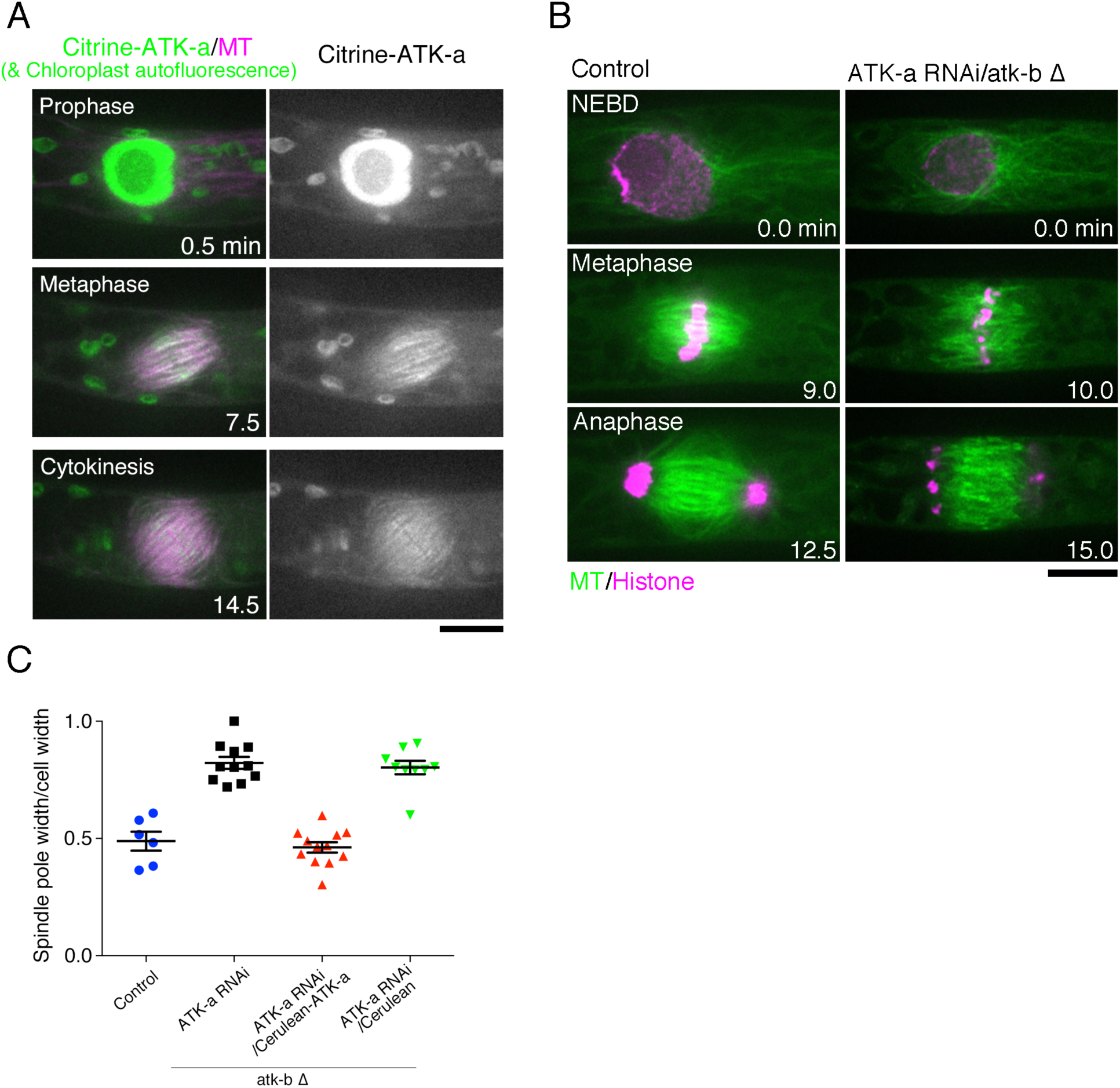
ATK/kinl4-I is required for spindle coalescence. **(A)** Localisation of Citrine-tagged ATK-a proteins. Bar, 10 μm. **(B)** Mitosis imaging of an ATK-a RNAi line where the paralogous ATK-b gene had been deleted. Bar, 10 μm. **(С)** Ratio of spindle width to cell width (±SE, n = 6 (control), n = 11 (ATK-a RNAi, n = 12 (ATK-a RNAi/Cerulean-ATK-a) n = 9 (ATK-a RNAi/Ceralean)), p < 0.0009 (Mann Whitney test)). The width at the spindle edge was measured.

**Table S1.**
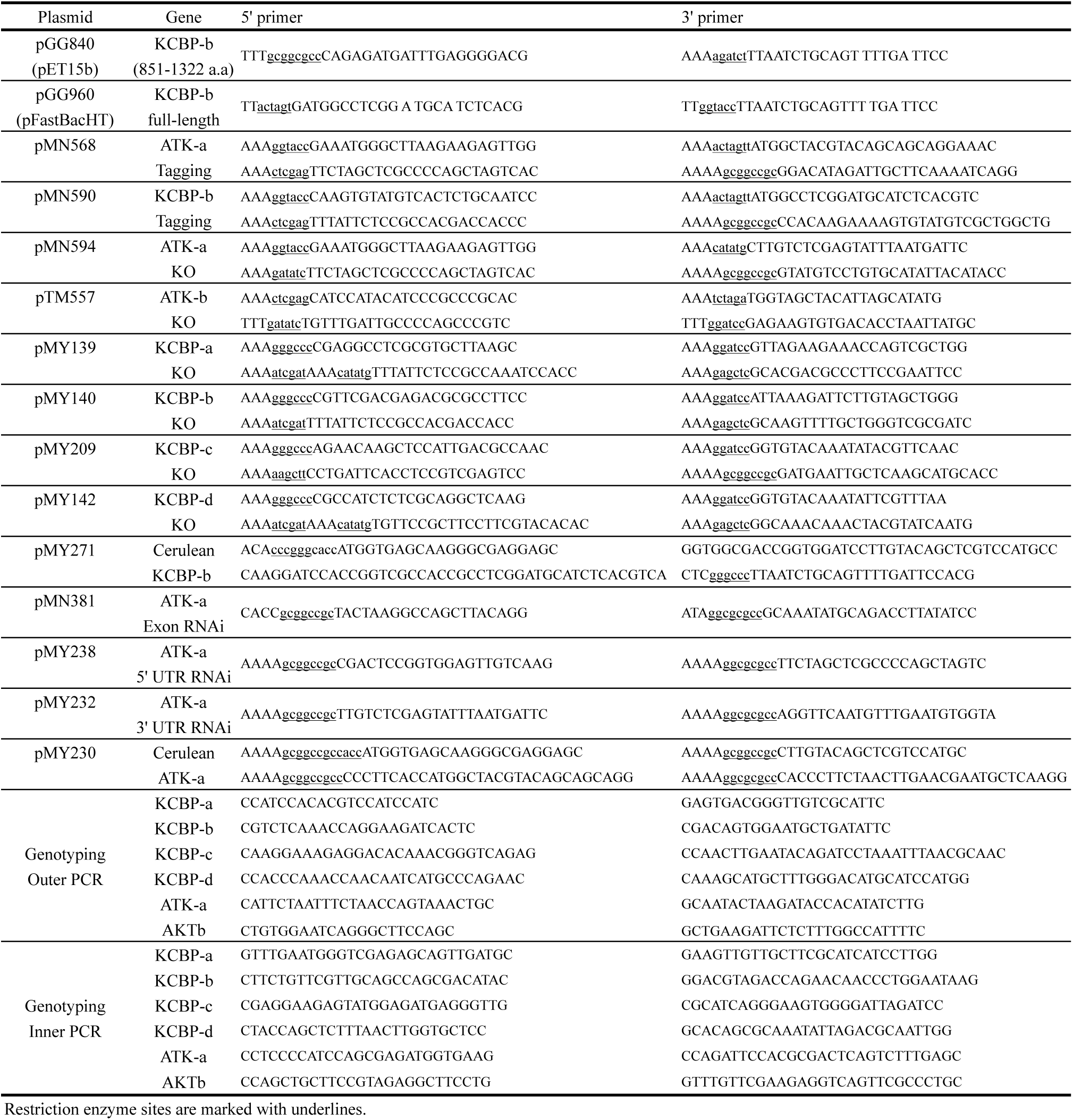
List of PCR primers used for plasmid construction. Restriction enzyme sites are marked with underlines.

**Table S2.** Moss lines used in this study. G418: Gentamicin, Zeo: Zeocin, Hyg: Hygromycin, BSD: Blasticidin S, NTC: Nourseothricin

| Line # | Clone # | Name | Plasmid | Resistance | References |
| --- | --- | --- | --- | --- | --- |
| 0002 | 3 | GFP-tubulin/histone H2B-mRFP | pGG616 | G418, Zeo | Nakaoka et al., 2012 |
| 0379 | 1 | EB1-Citrine/mCherry-tubulin | pKK37 | G418, Zeo | Kosetsu et al., 2013 |
| 0409 | 7,10 | ATK-b KO | pTM557 | Zeo,G418,NTC |  |
| 0414 | 34, 129 | mCherry-tubulin/Citrine-ATK-a | pMN568 | BSD |  |
| 0428 | 1-10 | ATK-b KO /ATK-a exon RNAi | pMN381 | Zeo, G418, NTC, Hyg |  |
| 0479 | 20, 21, 27 | KCBP-abcd KO | pMY209 | Zeo, G418, NTC, BSD |  |
| 0485 | 29, 86, 167 | mCherry-tubulin/Citrine-KCBP-b | pMN590 | BSD | Jonsson et al., 2015 |
| 0497 | 9, 20 | mCherry-tubulin/Citrine-ATK-a/ ATK-b KO | pTM557 | BSD |  |
| 0499 | 1-10 | ATK-b KO/ ATK-a 5'UTR RNAi | pMY238 | Zeo, G418, NTC, Hyg |  |
| 0500 | 1-10 | ATK-b KO/ ATK-a 3'UTR RNAi | pMY232 | Zeo, G418, NTC, Hyg |  |
| 0548 | 7, 8 | ATK-b KO/ ATK-a 5'UTR RNAi/Cerulean-ATK-a | pMY230 | Zeo, G418, NTC, Hyg, BSD |  |
| 0550 | 8, 11 | ATK-b KO/ ATK-a 5'UTR RNAi/Cerulean | pTM416 | Zeo, G418, NTC, Hyg, BSD |  |
| 0554 | 3, 105 | KCBP-abcd KO/EB1-mCherry | pMY295 | G418, Zeo, BSD |  |
| 0555 | 1, 4, 6, 8 | KCBP-abcd KO/Cerulean-KCBP-b | pMY271 | G418, Zeo, BSD |  |

